# Circ_0005962 functions as an oncogene to aggravate non-small cell lung cancer progression via circ_0005962/miR-382-5p/PDK4 regulatory network

**DOI:** 10.1101/2020.03.06.980185

**Authors:** Zhihong Zhang, Zhenxiu Shan, Rubin Chen, Xiaorong Peng, Bin Xu, Liang Xiao, Guofei Zhang

**Affiliations:** Department of Oncology, Gong’an County People’s Hospital, Hubei 433000, China; Department of Radiology, Gong’an County People’s Hospital, Hubei 433000, China; Department of Pathology, Gong’an County People’s Hospital, Hubei 433000, China; Department of Cerebral Surgery, Gong’an County People’s Hospital, Hubei 433000, China; Department of Gastrointestinal Surgery, Gong’an County People’s Hospital, Hubei 433000, China

**Keywords:** circ_0005962, miR-382-5p, PDK4, NSCLC

## Abstract

Non-small cell lung cancer (NSCLC) is a leading threat to human lives with high incidence and mortality. Circular RNAs (circRNAs) were reported to play important roles in human cancers. The purpose of this study was to investigate the role of circ_0005962 and explore the underlying functional mechanisms. The expression of circ_0005962, miR-382-5p and pyruvate dehydrogenase kinase 4 (PDK4) was detected by quantitative real-time polymerase chain reaction (qRT-PCR). Cell proliferation and cell apoptosis were assessed by cell counting kit-8 (CCK-8) assay and flow cytometry assay, respectively. The protein levels of Beclin 1, light chain3 (LC3-II/LC3-I), PDK4, Cleaved Caspase 3 (C-caspase 3) and proliferating cell nuclear antigen (PCNA) were examined using western blot analysis. Glycolysis was determined according to the levels of glucose consumption and lactate production. The interaction between miR-382-5p and circ_0005962 or PDK4 was predicted by the online tool CircInteractome or starbase and verified by dual-luciferase reporter assay and RNA immunoprecipitation (RIP) assay. Xenograft model was constructed to investigate the role of circ_0005962 *in vivo*. circ_0005962 expressed with a high level in NSCLC tissues and cells. Circ_0005962 knockdown inhibited proliferation, autophagy, and glycolysis but promoted apoptosis in NSCLC cells. MiR-382-5p was targeted by circ_0005962, and its inhibition reversed the role of circ_0005962 knockdown. Besides, PDK4, a target of miR-382-5p, was regulated by circ_0005962 through miR-382-5p, and its overexpression abolished the effects of miR-382-5p reintroduction. Circ_0005962 knockdown suppressed tumor growth *in vivo*. Circ_0005962 knockdown restrained cell proliferation, autophagy, and glycolysis but stimulated apoptosis through modulating the circ_0005962/miR-382-5p/PDK4 axis. Our study broadened the insights into understanding the mechanism of NSCLC progression.

## Introduction

Lung cancer is a leading cause of cancer-related death worldwide (Magnuson et al., 2016). Lung cancer is the second most common cancer among men and women: second only to prostate cancer in men, and second only to breast cancer in women (Testa et al., 2018). Non-small cell lung cancer (NSCLC) and small cell lung cancer are two types of lung cancer, and NSCLC accounts for about 85% of all lung cancer cases (Zhukovsky et al., 2014). NSCLC, including large cell carcinoma, squamous cell carcinoma, and adenocarcinoma, is associated with high incidence and mortality (Barnett et al., 2016; Brody, 2014). Clinically, treatment modalities, including chemotherapy and surgery, are used to treat NSCLC, but the 5-year overall survival rate for all stages of NSCLC patients is only 16% (Laskin et al., 2005; Testa et al., 2018). The severe situation of NSCLC treatment makes it urgent to further explore the mechanism of occurrence and development of NSCLC to establish novel therapeutic strategies.

Circular RNAs (circRNAs) belong to non-coding RNAs and derive from alternative and back-splicing of precursor mRNAs. CircRNAs are mostly detected in the cytoplasm and function as competing endogenous RNAs (ceRNAs) to work as microRNA (miRNA) sponges (Hansen et al., 2013; Kulcheski et al., 2016; Salzman et al., 2012). Advances of high-throughput RNA sequencing in the identification of circRNAs hinted that circRNAs participated in the pathogenesis of cancers (Liang et al., 2014). Recent studies stated that circRNAs played vital roles in the development of NSCLC. For example, circ_100146 functioned as an oncogene, and its suppression hindered cell proliferation and invasion (L. Chen et al., 2019). CircP4HB showed a high level in NSCLC, promoted epithelial-mesenchymal transition (EMT), and was associated with metastatic diseases (T. Wang et al., 2019). CircRNA F-circEA-2a contributed to cell migration and invasion but had little role in cell proliferation (Tan et al., 2018). A former study obtained dozens of circRNAs through the CircBase database and CSCD database for comparison between lung adenocarcinoma tissues and paired non-tumor tissues (Liu et al., 2019), and circ_0005962, back-spliced from tyrosine 3-monooxygenase/tryptophan 5-monooxygenase activation protein zeta (YWHAZ), was one of the significantly upregulated circRNAs and had valuable prognostic significance. However, the understanding of the specific roles of circ_0005962 in NSCLC remains lacking and requires further exploration.

It is well documented that miRNAs are involved in the biological processes of cancers. MiRNAs are known as small non-coding RNAs with 18∼23 nucleotides (Jiang et al., 2014). MiRNAs play vital roles in tumor formation, growth and metastasis by acting as oncogenes or tumor suppressor genes. Among these miRNAs, miR-382-5p was frequently mentioned in different cancers, including glioma, oral squamous cell carcinoma and breast cancer (Ho et al., 2017; Sun et al., 2019; J. Wang, C. Chen, et al., 2019). Unfortunately, the role of miR-382-5p in NSCLC is unknown, and we attempted to investigate the function of miR-382-5p in NSCLC cells.

One of the important functions of pyruvate dehydrogenase kinase (PDK) is to regulate the metabolic conversion from mitochondrial respiration to cytoplasmic glycolysis (Jeoung, 2015). Glycolysis is a preferential way for tumor cells to obtain energy (Vander Heiden et al., 2009). Pyruvate dehydrogenase kinase 4 (PDK4) is one of the PDK family protein kinases and located on chromosome 7q21.3 (J. Wang, Y. Qian, et al., 2019). PDK4 was reported to be implicated in numerous cellular activities, such as cell proliferation, metastasis, drug resistance, glycolysis and autophagy in different types of cancers (Feng et al., 2019; J. Wang, Y. Qian, et al., 2019; Yang et al., 2019). The specific role of PDK4 and associated action mechanisms in NSCLC still need to be explored to expand the function of PDK4 in cancers.

In the present study, we measured the expression of circ_0005962 in NSCLC tissues and cells. Functional analyses revealed the role of circ_0005962 in cell proliferation, autophagy, glycolysis and apoptosis. Besides, we constructed circRNA-miRNA-mRNA regulatory axis to expound the potential mechanism of circ_0005962 in NSCLC. Our study aimed to provide new sights for the understanding of NSCLC development and supply promising biomarkers.

## Materials and methods

### Tissues and cell lines

A total of 45 tumor tissues and adjacent normal tissues from NSCLC patients were collected from Gong’an County People’s Hospital. Informed consent was signed by each subject. All tissues were immediately placed into liquid nitrogen and stored at −80□ ultra low-temperature refrigerator. This research obtained the approval of the Ethics Committee of Gong’an County People’s Hospital.

NSCLC cell lines (A549 and HCC827) and human bronchial epithelial cells (BEAS-2B) were purchased from Zishi Biotechnology (Shanghai, China). A549 and HCC827 cells were maintained in 90% Roswell Park Memorial Institute 1640 (RPMI 1640; Sigma, St. Louis, MO, USA) containing 10% fetal bovine serum (FBS; Sigma). BEAS-2B cells were cultured in 90% Dulbecco’s Modified Eagle Medium (DMEM; Sigma) containing 10% FBS (Sigma). All mediums were placed at 37□ containing 5% CO_2_.

### Quantitative real-time polymerase chain reaction (qRT-PCR)

Trizol reagent (Beyotime, Shanghai, China) was utilized to isolate total RNA from tissues and cells. The complementary DNA (cDNA) was assembled using the riboSCRIPT Reverse Transcription Kit (Ribobio, Guangzhou, China). Then the amplification reaction was carried out using SYBR Green Master PCR mix (Beyotime) through the ABI 7900 system (Applied Biosystems, Foster City, CA, USA). The relative expression was normalized by Glyceraldehyde-3-phosphate dehydrogenase (GAPDH) or small nuclear RNA U6 and calculated using 2^−ΔΔCt^ method. The primers used were listed as below: circ_0005962, forward (F): 5’-AACTCCCCAGAGAAAGCCTGC-3’ and reverse (R): 5’-TGCTTGTGAAGCATTGGGGAT-3’; YWHAZ, forward (F): 5’-ACTTTTGGTACA TTGTGGCTTCAA-3’ and reverse (R): 5’-CCGCCAGGACAAACCAGTAT-3’; PDK4, F: 5’-GGAGCATTTCTCGCGCTACA-3’ and R: 5’-ACAGGCAATTCTTGTCGCAAA-3’; GAPDH, F: 5’-CTGGGCTACACTGAGCACC-3’ and R: 5’-AAGTGGTCGTTGAGGGCAATG-3’; miR-382-5p, F: 5’-ATCCGTGAAGTTGTTCGTGG-3’ and R: 5’-TATGGTTGTAGAGGACTCCTTGAC-3’; U6, F: 5’-GCUUCGGCAGCACAUAUACUAAAAU-3’ and R: 5’-CGCUUCACGAAUUUGCGUGUCAU-3’.

### RNase R treatment

To confirm the stability and tolerance of circ_0005962, the RNA extraction was probed with or without RNase R (Applied Biological Materials Inc., Vancouver, Canada) at 37□ for 10 min. Then, the qRT-PCR analysis was conducted as described above.

### Cell transfection

Small interference RNA against circ_0005962 and negative control were synthesized by Sangon Biotech (Shanghai, China). The mimics of miR-382-5p (miR-382-5p), the inhibitor of miR-382-5p (anti-miR-382-5p) and respective negative control (NC or anti-NC) were purchased from Ribobio (Shanghai, China). The overexpression of circ_0005962 (circ_0005962) was performed as the previous study described (Y. Wang, J. Zhang, et al., 2019) and constructed by Genechem (Shanghai, China), while the specific plasmid without the circ_0005962 cDNA was served as control (circ-NC). The vector pcDNA3.1-PDK4 (PDK4) for the overexpression of PDK4 and the control pcDNA empty vector (vector) were constructed by Sangon Biotech. Lentiviral vector (Lenti-short hairpin) for stable NEAT1 downregulation (Lv-sh-circ_0005962) and corresponding negative control (Lv-sh-NC) were obtained from Genechem. Cell transfection was conducted by using Lipofectamine 3000 (Invitrogen, Carlsbad, CA, USA). Transfection efficiency and following experiments were implemented after 48h of transfection.

### Cell counting kit-8 (CCK-8) assay

Cell proliferation was investigated by CCK-8 assay. A549 or HCC827 cells with different transfection were seeded into 96-well plates (5×10^3^ cells/well). Then, cells were interacted with CCK-8 solution (Beyotime) for continuing 2 h at 24, 48, and 72 h. The absorbance at 450 nm was measured using a microplate reader (Bio-Rad, Hercules, CA, USA).

### Flow cytometry assay

Flow cytometry was applied to monitor cell apoptosis. After 48 h, A549 or HCC827 cells with different transfection were washed with phosphate buffer saline (PBS), probed with 0.25% trypsin and resuspended in binding buffer. Afterward, the Annexin V-fluorescein isothiocyanate (FITC)/propidium iodide (PI) apoptosis detection kit (Sigma) was used to doubly stain cells for 15 min in the dark. Eventually, the apoptotic cells were sorted using CellQuest software under a flow cytometer (Becton Dickinson, Franklin Lakes, NJ, USA).

### Western blot

The western blot analysis was performed in line with a previous study described (W. Ren et al., 2019). In brief, total proteins were separated and transferred into polyvinylidene fluoride (PVDF) membranes (Bio-Rad). After the block, the membranes were incubated with the primary antibodies and the secondary antibodies. Finally, the seeking proteins were visualized using the enhanced chemiluminescent reagent (Beyotime) through an imaging system (Bio-Rad). The antibodies used were listed as follows: anti-Beclin 1 (1:1000; cat. no. ab210498; Abcam, Cambridge, MA, USA), anti-light chain3 (LC3) (1:2000; cat. no. ab192890), anti-PDK4 (1:1000; cat. no. ab89295), anti-Cleaved Caspase 3 (C-caspase 3) (1:1000; cat. no. ab2302), anti-proliferating cell nuclear antigen (PCNA) (1:1000; cat. no. ab92552), anti-GAPDH (1:1000; cat. no. ab8245) and the horseradish peroxidase-conjugated secondary antibodies (1:5000; cat. no. ab205718).

### The detection of glucose consumption and lactate production

A549 or HCC827 cells with different transfection were planted into 96-well plates. After 48 h, cells were collected, washed three times with PBS, and then used for the detection of glucose consumption and lactate production using the Glucose Uptake Colorimetric Assay Kit and Lactate Assay Kit (Sigma) in agreement with the manufacturer’s instructions.

### Bioinformatics analysis

The targets of circRNA and miRNA were forecasted, and their binding sites were analyzed by the online bioinformatics tool CircInteractome (https://circinteractome.nia.nih.gov/) and starbase (http://starbase.sysu.edu.cn/).

### Dual-luciferase reporter assay

Dual-luciferase reporter assay was carried out for the verification of the relationship between miR-382-5p and circ_0005962 or PDK4. The partial frame of circ_0005962 containing the binding site or mutant binding site (wild-type or mutant-type) with miR-382-5p was amplified and cloned into the downstream of pGL4 vector (Promega, Madison, WI, USA) to generate circ_0005962-wt and circ_0005962-mut fusion plasmids. Likewise, the 3’ UTR sequences of PDK4 harboring the binding site with miR-382-5p or mutant binding site (wild-type or mutant-type) were also amplified and inserted into the downstream of pGL4 vector to generate PDK4-wt and PDK-mut fusion plasmids. Subsequently, these fusion plasmids and miR-382-5p or NC were co-transfected into A549 and HCC827 cells, respectively. After 48 h, the cells were collected and detected using the Dual-luciferase assay system (Promega). The firefly luciferase activity was normalized by Renilla luciferase activity.

### RNA immunoprecipitation (RIP) assay

RIP assay was executed to further confirm the relationship between miR-382-5p and circ_0005962 or PDK4. In one case, A549 and HCC827 cells were harvested and incubated with RNA lysis buffer. Then, cell lysate was incubated with RIP binding buffer containing magnetic beads coated with human Ago2 antibody or mouse IgG antibody (control) (Millipore, Billerica, MA, USA). Subsequently, the levels of circ_0005962 and miR-382-5p enriched in the beads were detected by qRT-PCR. In another case, A549 and HCC827 cells transfected with miR-382-5p were lysed in RIP buffer with anti-Ago2- or IgG-bound magnetic beads. Next, the mRNA level of PDK4 enriched in the beads was examined by qRT-PCR.

### *In vivo* experiment

The mice experiment obtained the approval of the Animal Care and Use Committee of Gong’an County People’s Hospital. A total of 10 BALB/c nude mice (five-week-old, male) were purchased from HFK Bioscience (Beijing, China). A549 cells with Lv-sh-circ_0005962 or Lv-sh-NC transfection were subcutaneously injected into the back right flank of nude mice, dividing into the Lv-sh-circ_0005962 group and Lv-sh-NC group. Tumor volume was observed and recorded once a week following the algorithm (length×width^2^×0.5), lasting 5 weeks. In the end, the mice were killed, and tumor samples were removed for weighting and further molecular studies.

### Data analysis

The data were obtained from at least 3 times independent experiments analysis and conducted using SPSS 18.0 software (SPSS, Inc., Chicago, IL, USA). The survival curve was depicted via the Kaplan-Meier method. The correlation analysis was performed based on Spearman’s correlation coefficient. The differences between 2 groups were analyzed by Student’s *t*-test or one-way analysis of variance among multiple groups. The data after processing were presented as the mean ± standard deviation (SD), and *P* < 0.05 was considered to be statistically significant.

## Result

### High expression of circ_0005962 was observed in NSCLC tissues and cells and predicted the low survival rate of NSCLC patients

The expression of circ_0005962 was detected in NSCLC tissues and cell lines to observe whether circ_0005962 aberrantly regulated in NSCLC. As shown in Figure 1A, the expression of circ_0005962 was significantly higher in tumor tissues (n=45) than that in adjacent normal tissues (n=45). Besides, the expression of circ_0005962 was abundant in A549 and HCC827 cells relative to BEAS-2B cells (Figure 1B). Moreover, the survival curve was depicted utilizing Kaplan-Meier survival rate analysis according to the living status of NSCLC patients, and we found that the survival rate of patients with high expression of circ_0005962 was notably weaker than patients with low circ_0005962 expression (Figure 1C). Additionally, the result of qRT-PCR showed that the expression of circ_0005962 in RNase R+ group decreased a little compared with that in RNase R-group, while the expression of the parental linear mRNA (YWHAZ) was significantly reduced with the treatment of RNase R, indicating that circ_0005962 was resistant to RNase R (Figure 1D). The data indicated that circ_0005962 was aberrantly upregulated in NSCLC.

**Figure 1.**
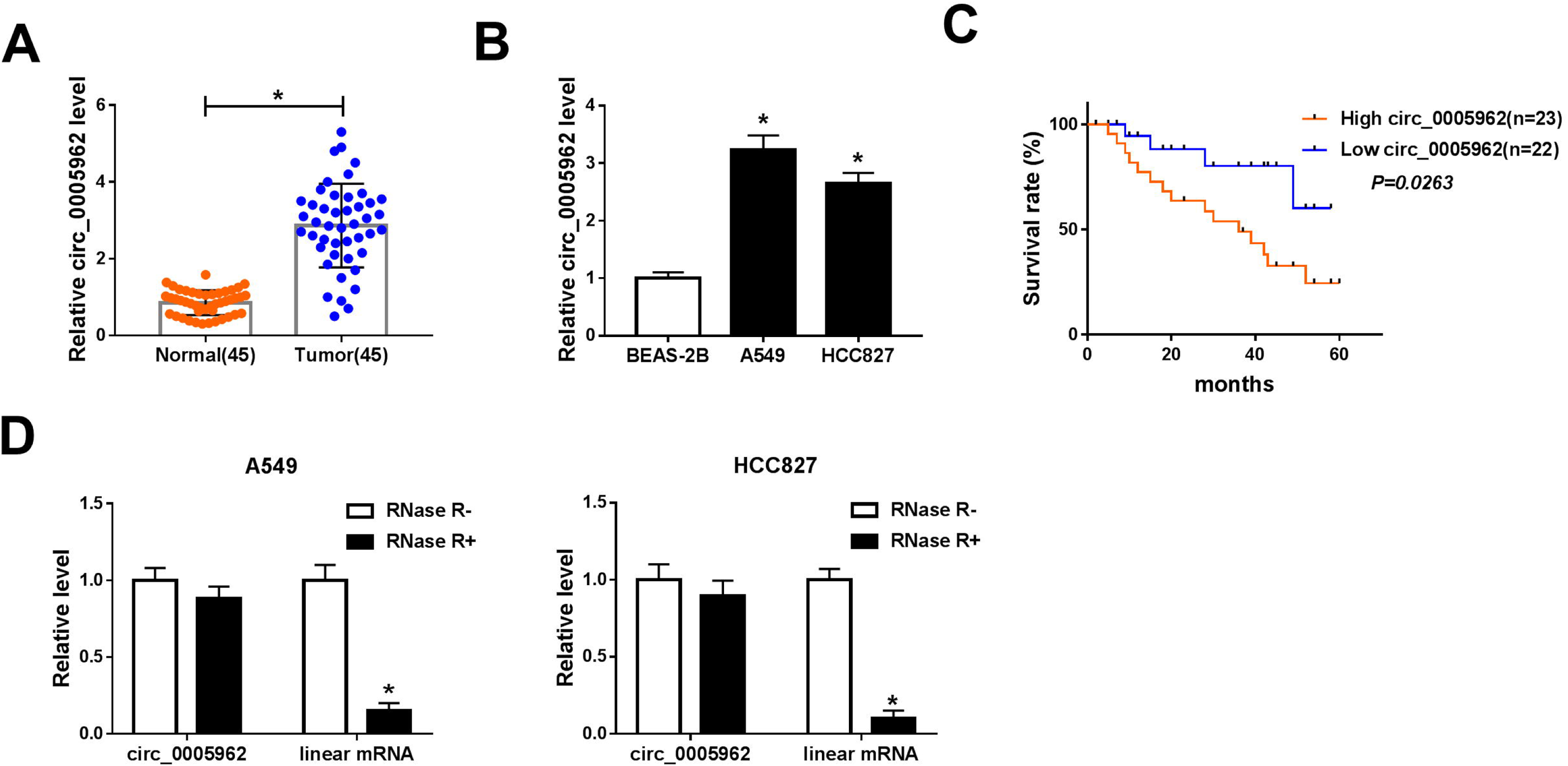
Circ_0005962 was highly expressed in NSCLC tissues and cells. (A) The expression of circ_0005962 in tumor tissues (n=45) and adjacent normal tissues (n=45) was detected by qRT-PCR. (B) The expression of circ_0005962 in NSCLC cell lines (A549 and HCC827) and normal cells (BEAS-2B) was detected by qRT-PCR. (C) The survival curve was depicted according to the Kaplan-Meier plot and analyzed by the log-rank test. (D) The tolerance of circ_0005962 and corresponding linear RNA to RNase was measured according to their mRNA levels. \**P* < 0.05.

### Circ_0005962 knockdown inhibited proliferation, autophagy, and glycolysis but promoted apoptosis in NSCLC cells

The endogenous level of circ_0005962 was knocked down to investigate the role of circ_0005962 in NSCLC cells. Si-circ_0005962 was inserted into the mature sequence of circ_0005962 to reduce circ_0005962 expression, si-NC acting as a control (Figure 2A). Then, the knockdown efficiency was examined by qRT-PCR, and the result showed that the expression of circ_0005962 in A549 and HCC827 cells with si-circ_0005962#1, si-circ_0005962#2 and si-circ_0005962#3 transfection was notably decreased than that with si-NC transfection, and the knockdown efficiency in the si-circ_0005962#1 group was the highest (Figure 2B). Hence, si-circ_0005962#1 was chosen for the following analyses. The result of CCK-8 assay revealed that the cell proliferation was prominently suppressed in A549 and HCC827 cells transfected with si-circ_0005962#1 (Figure 2C). On the contrary, flow cytometry assay presented that the apoptosis rate in A549 and HCC827 cells with si-circ_0005962#1 transfection was inversely enhanced (Figure 2D). Besides, the protein levels of autophagy-related markers were quantified to assess the change of autophagy, and we found that the levels of Beclin 1 and LC3-II/LC3-I were declined with circ_0005962 knockdown (Figure 2E and 2F). Moreover, the levels of glucose in culture medium and lactate production were checked to assess glycolysis progression, and we noticed that the level of existing glucose in A549 and HCC827 cells with si-circ_0005962#1 transfection was higher than that with si-NC transfection, suggesting that the consumptive glucose was reduced (Figure 2G). The level of lactate production was significantly blocked in A549 and HCC827 cells with si-circ_0005962#1 transfection relative to si-NC (Figure 2H). All data clarified that circ_0005962 knockdown inhibited proliferation, autophagy, and glycolysis but induced apoptosis in NSCLC cells.

**Figure 2.**
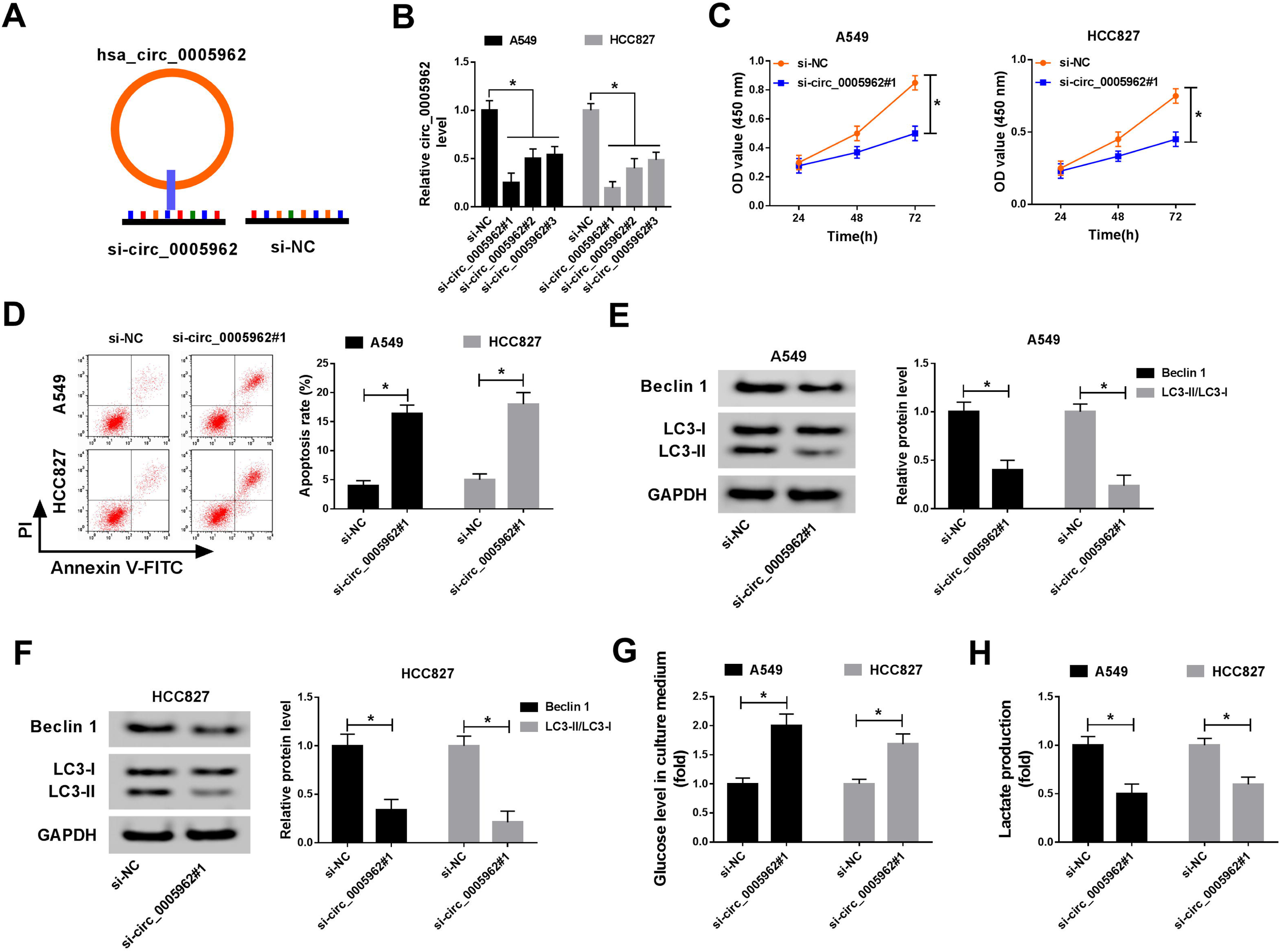
Circ_0005962 knockdown inhibited proliferation, autophagy, and glycolysis but contributed to apoptosis in NSCLC cells. (A) The diagram of circ_0005962 knockdown. (B) The efficiency of circ_0005962 knockdown in different transfection lines was detected by qRT-PCR. A549 and HCC827 cells were transfected with si-circ_0005962#1 or si-NC. (C) Cell proliferation was assessed by CCK-8 assay. (D) Cell apoptosis was executed by flow cytometry assay. (E and F) The protein levels of Beclin 1 and LC3-II/LC3-I were quantified by western blot. (G and H) The glycolysis progression was evaluated according to the level of glucose in culture medium and lactate production. \**P* < 0.05.

### MiR-382-5p was a target of circ_0005962

Generally, circ_0005962 functioned by acting as a ceRNA to modulate the expression of target miRNAs. The putative target miRNAs were predicted by the online tool CircInteractome, and miR-382-5p was one of target miRNAs of circ_0005962 with specific binding sites (Figure 3A). To ascertain the relationship between circ_0005962 and miR-382-5p, dual-luciferase reporter assay and RIP assay were performed. The luciferase activity in A549 and HCC827 cells with circ_0005962-wt and miR-382-5p transfection was substantially decreased compared with circ_0005962-wt and NC transfection, while the luciferase activity in A549 and HCC827 cells with circ_0005962-mut and miR-382-5p transfection was no difference compared with circ_0005962-mut and NC transfection (Figure 3B). The RIP analysis detected the higher expression of circ_0005962 and miR-382-5p in the Ago2 pellet of A549 and HCC827 lysate than that in the IgG control (Figure 3C). Moreover, the qRT-PCR analysis exhibited that the expression of miR-382-5p was notably declined with circ_0005962 overexpression but improved with circ_0005962 knockdown in A549 and HCC827 cells (Figure 3D). Additionally, the expression of miR-382-5p in NSCLC tumor tissues (n=45) was markedly weaker than that in normal tissues (n=45) (Figure 3E). The expression of miR-382-5p in A549 and HCC827 cells was consistently lower than that in BEAS-2B cells (Figure 3F). Spearman’s correlation coefficient revealed that circ_0005962 expression was negatively correlated with miR-382-5p expression in NSCLC tissues (Figure 3G). These analyses maintained that miR-382-5p was a target of circ_0005962, and its expression was regulated by circ_0005962.

**Figure 3.**
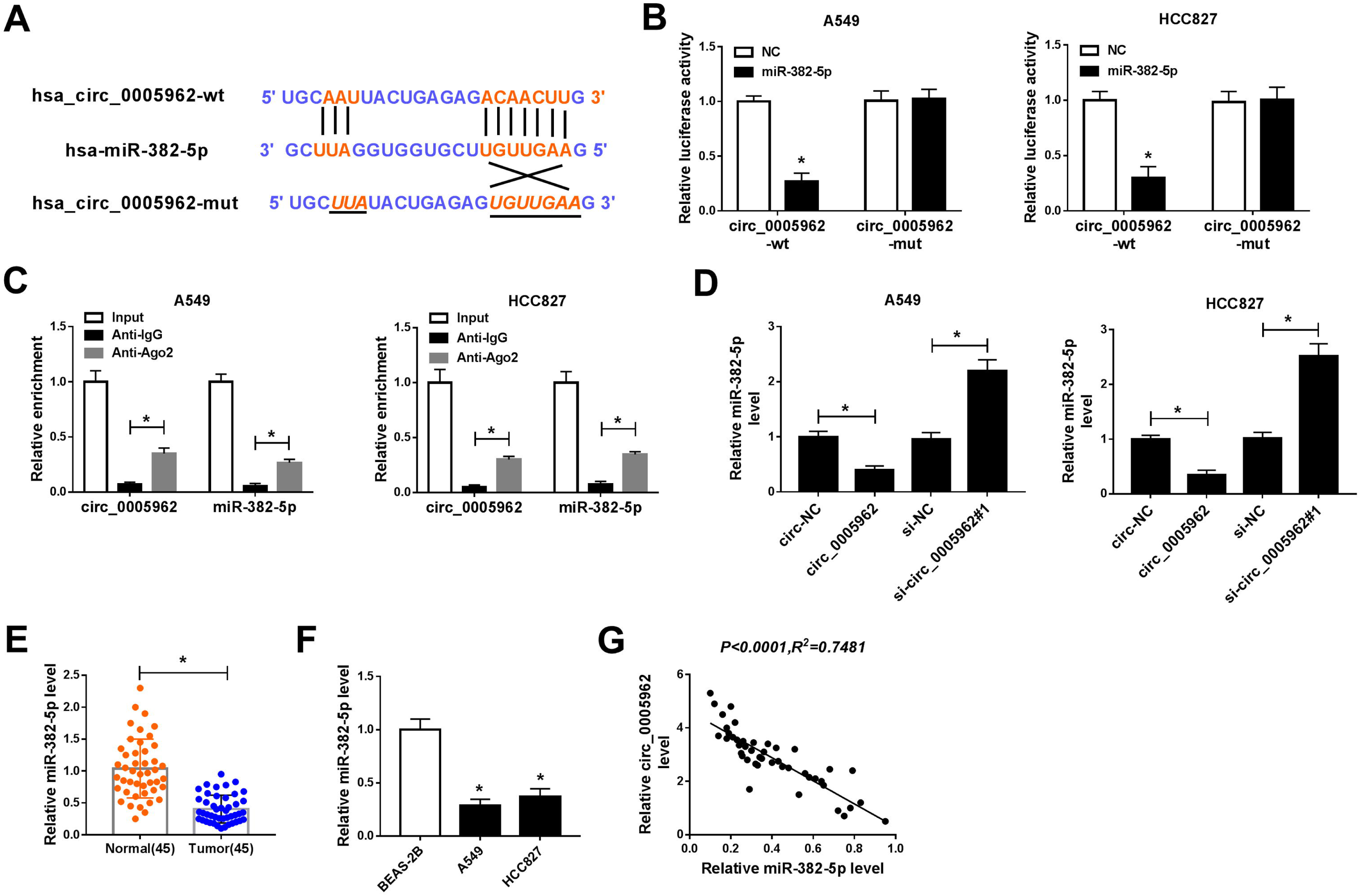
MiR-382-5p was a target of circ_0005962. (A) The binding sites between circ_0005962 and miR-382-5p were analyzed by the online tool CircInteractome. (B) The relationship between circ_0005962 and miR-382-5p was verified by dual-luciferase reporter assay. (C) The relationship between circ_0005962 and miR-382-5p was further confirmed by RIP assay. (D) The expression of miR-382-5p in A549 and HCC827 cells transfected with circ_0005962 or si-circ_0005962 was detected by qRT-PCR. (E and F) The expression of miR-382-5p in NSCLC tissues and cell lines was detected by qRT-PCR. (G) The correlation between circ_0005962 and miR-382-5p was analyzed by Spearman’s correlation coefficient. \**P* < 0.05.

### Inhibition of miR-382-5p reversed the role of circ_0005962 knockdown in NSCLC cells

A549 and HCC827 cells were introduced with si-circ_0005962#1, si-NC, si-circ_0005962#1+anti-miR-382-5p and si-circ_0005962#1+anti-NC, respectively. First, the expression of miR-382-5p in these transfected cells was checked, and we found that the expression of miR-382-5p was enhanced in the si-circ_0005962#1 but inhibited in the si-circ_0005962#1+anti-miR-382-5p group (Figure 4A). The cell proliferation inhibited by si-circ_0005962#1 was recovered in A549 and HCC827 cells with si-circ_0005962#1+anti-miR-382-5p transfection (Figure 4B). The elevated cell apoptosis rate in cells with si-circ_0005962#1 transfection was suppressed in cells with si-circ_0005962#1+anti-miR-382-5p transfection (Figure 4C). The protein levels of Beclin 1 and LC3-II/LC3-I were depleted in A549 and HCC827 cells transfected with si-circ_0005962#1 but restored in cells transfected with si-circ_0005962#1+anti-miR-382-5p (Figure 4D and 4E). The level of existing glucose in medium in the si-circ_0005962#1+anti-miR-382-5p group was declined than that in the si-circ_0005962#1 group (Figure 4F). The level of lactate production inhibited in the si-circ_0005962#1 group was promoted in the si-circ_0005962#1+anti-miR-382-5p group (Figure 4G). These results meant that circ_0005962 knockdown inhibited proliferation, autophagy, and glycolysis but induced apoptosis through enhancing the expression of miR-382-5p.

**Figure 4.**
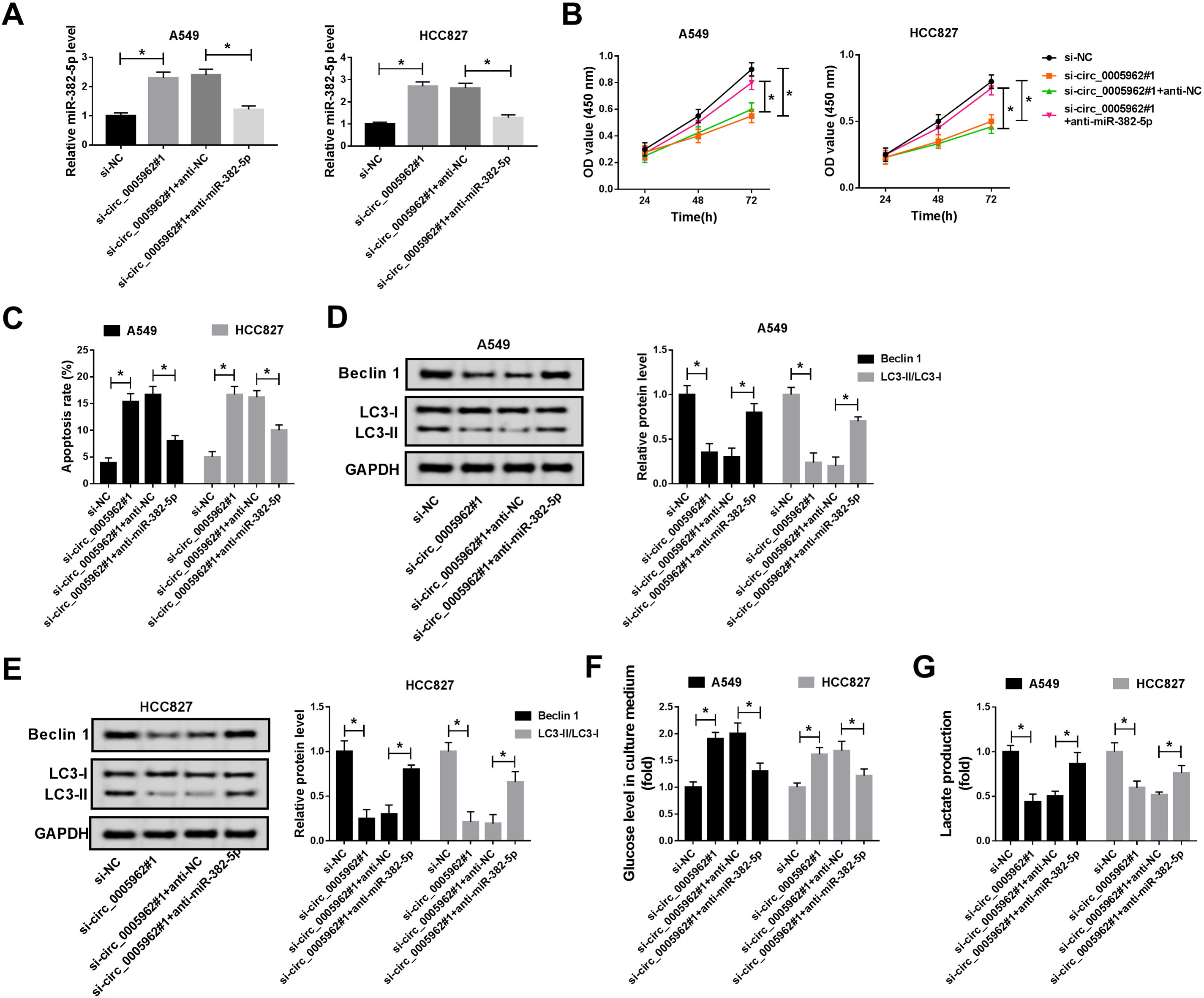
Inhibition of miR-382-5p reversed the role of circ_0005962 knockdown. A549 and HCC827 cells were introduced with si-circ_0005962#1, si-NC, si-circ_0005962#1+anti-miR-382-5p or si-circ_0005962#1+anti-NC. (A) The transfection efficiency was examined by qRT-PCR. (B) Cell proliferation was assessed by CCK-8 assay. (C) Cell apoptosis was executed by flow cytometry assay. (D and E) The protein levels of Beclin 1 and LC3-II/LC3-I were quantified by western blot. (F and G) The glycolysis progression was evaluated according to the levels of glucose in culture medium and lactate production. \**P* < 0.05.

### PDK4 was a target of miR-382-5p

CircRNA-miRNA-mRNA regulatory network is an important mechanism participating in the development of human cancers. The target mRNAs of miR-382-5p were analyzed to observe that whether circ_0005962 functioned following this mechanism. Online bioinformatics tool starbase predicted that PDK4 was one of targets of miR-382-5p with a specific binding site at its 3’ UTR (Figure 5A). Besides, miR-382-5p overexpression predominantly inhibited the luciferase activity in A549 and HCC827 cells transfected with PDK4-wt but not PDK4-mut (Figure 5B). Moreover, the enrichment of PDK4 was notably elevated in the miR-382-5p-transfected group compared with that in the NC group after Ago2 RIP, while enrichment of PDK4 after IgG RIP showed no efficacy (Figure 5C). Next, the western blot analysis monitored that the level of PDK4 was weakened in A549 and HCC827 cells with miR-382-5p transfection but elevated in cells with circ_0005962+miR-382-5p transfection (Figure 5D). Also, the expression of PDK4 was detected in NSCLC tumor tissues, and the result presented that the expression of PDK4 at both mRNA and protein levels was abnormally higher in NSCLC tumor tissues (n=45) relative to normal tissues (n=45) (Figure 5E and 5F). Likewise, the protein level of PDK4 in A549 and HCC827 cells was also enhanced compared to BEAS-2B cells (Figure 5G). Furthermore, the mRNA level of PDK4 was positively correlated with circ_0005962 level but negatively correlated with the miR-382-5p level in NSCLC tumor tissues (Figure 5H and 5I). All data suggested that PDK4 was a target of miR-382-5p and regulated by miR-382-5p and circ_0005962.

**Figure 5.**
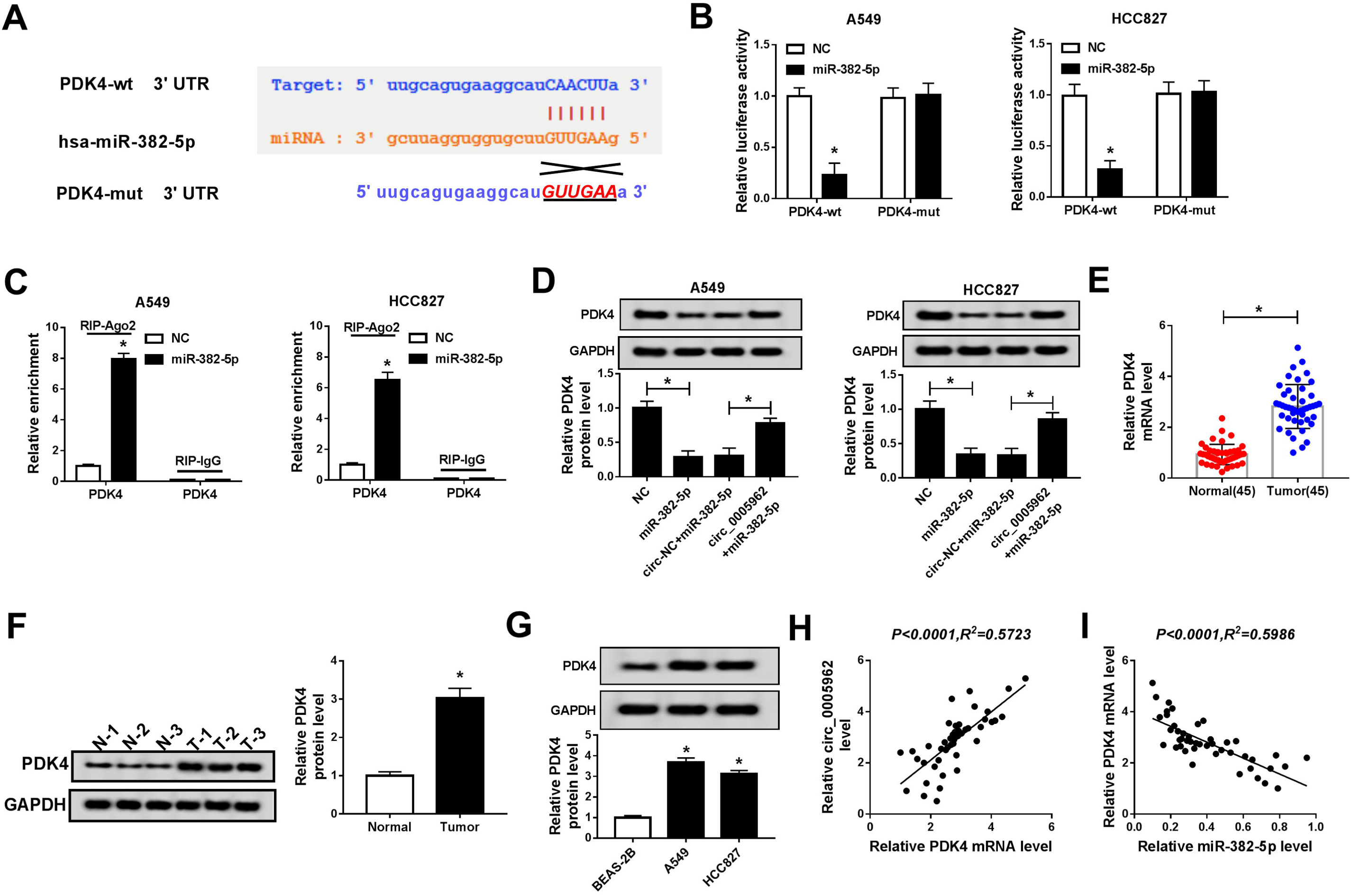
PDK4 was a target of miR-382-5p. (A) The binding sites between PDK4 3’ UTR and miR-382-5p were analyzed by online tool starbase. (B) The relationship between PDK4 and miR-382-5p was verified by dual-luciferase reporter assay. (C) The relationship between PDK4 and miR-382-5p was further confirmed by RIP assay. (D) The expression of PDK4 at the protein level in A549 and HCC827 cells transfected with miR-382-5p or circ_0005962+miR-382-5p was detected by western blot. (E and F) The expression of PDK4 at mRNA and protein levels in NSCLC tissues was detected by qRT-PCR and western blot. (G) The protein level of PDK4 in NSCLC cell lines was detected by western blot. (H and I) The correlation between PDK4 and circ_0005962 or miR-382-5p was analyzed by Spearman’s correlation coefficient. \**P* < 0.05.

### PDK4 overexpression abolished the role of miR-382-5p reintroduction in NSCLC cells

A549 and HCC827 cells were introduced with miR-382-5p, NC, miR-382-5p+PDK4 and miR-382-5p+vector, respectively. The expression of PDK4 was examined to assess transfection efficiency, and the result showed that the level of PDK4 was obviously reduced in cells transfected with miR-382-5p but regained in cells transfected with miR-382-5p+PDK4 (Figure 6A). Afterwards, the cell proliferation was inhibited by miR-382-5p reintroduction but promoted by the combination of miR-382-5p reintroduction and PDK4 overexpression (Figure 6B). The apoptosis rate was elevated by miR-382-5p reintroduction but restrained by the combination of miR-382-5p reintroduction and PDK4 overexpression (Figure 6C). Moreover, the levels of Beclin1 and LC3-II/LC3-I were weakened in A549 and HCC827 cells transfected with miR-382-5p but rescued in cells transfected with miR-382-5p+PDK4 (Figure 6D and 6E). The existing glucose level was abundant in the miR-382-5p group but reduced in the miR-382-5p+PDK4 group, suggesting that PDK4 overexpression enhanced the level of glucose consumption inhibited by miR-382-5p reintroduction (Figure 6F). The level of lactate production was blocked by miR-382-5p reintroduction but recovered by the combination of miR-382-5p reintroduction and PDK4 overexpression (Figure 6G). These data hinted that miR-382-5p attenuated cell proliferation, autophagy, and glycolysis but contributed to apoptosis through inhibiting the expression of PDK4.

**Figure 6.**
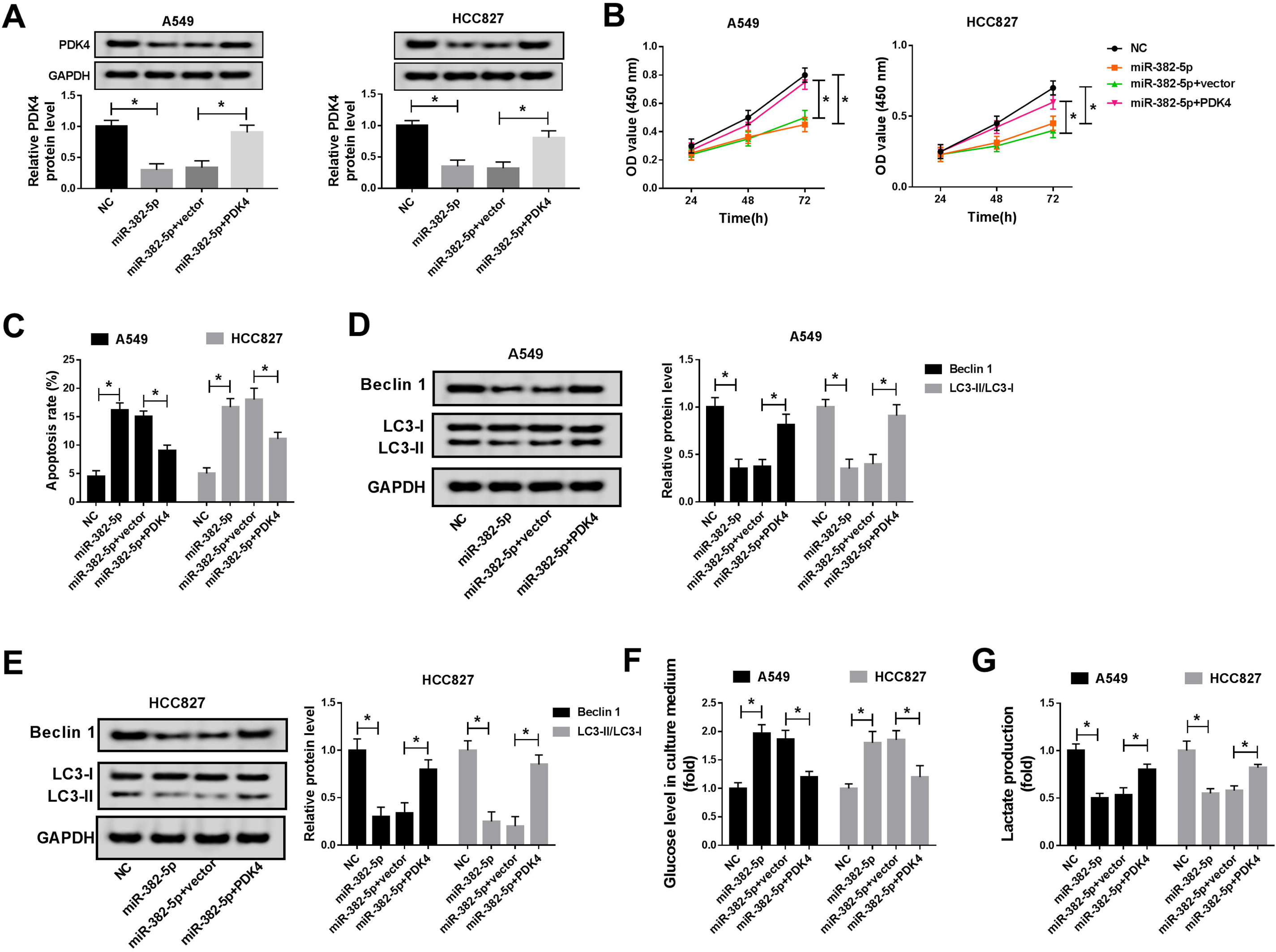
PDK4 overexpression abrogated the role of miR-382-5p reintroduction. A549 and HCC827 cells were transfected with miR-382-5p, NC, miR-382-5p+PDK4 or miR-382-5p+vector. (A) The transfection efficiency was checked according to the expression level of PDK4 using western blot. (B) Cell proliferation was assessed by CCK-8 assay. (C) Cell apoptosis was monitored by flow cytometry assay. (D and E) The protein levels of Beclin 1 and LC3-II/LC3-I were quantified by western blot. (F and G) The glycolysis progression was evaluated according to the level of glucose in culture medium and lactate production. \**P* < 0.05.

### Circ_0005962 knockdown inhibited tumor growth *in vivo*

A549 cells with stable Lv-sh-circ_0005962 transfection were subcutaneously injected into the groin of nude mice to determine the role of circ_0005962 *in vivo*. As shown in Figure 7A and 7B, circ_0005962 knockdown remarkably reduced the tumor volume and tumor weight. After injection for 5 weeks, all mice were killed, and the tumors were removed for expression analysis. The expression of circ_0005962 and the mRNA level of PDK4 were noticeably declined in the Lv-sh-circ_0005962 group, while the expression of miR-382-5p was conspicuously strengthened in the Lv-sh-circ_0005962 group (Figure 7C). Besides, the protein level of PDK4 was consistent with its mRNA level. Additionally, the levels of apoptosis-related marker (C-caspase 3) and proliferation-related marker (PCNA) were monitored, and we discovered that the level of C-caspase 3 was reinforced, while the level of PCNA plummeted in the Lv-sh-circ_0005962 group (Figure 7D). Collectively, circ_0005962 knockdown impeded tumor growth *in vivo*.

**Figure 7.**
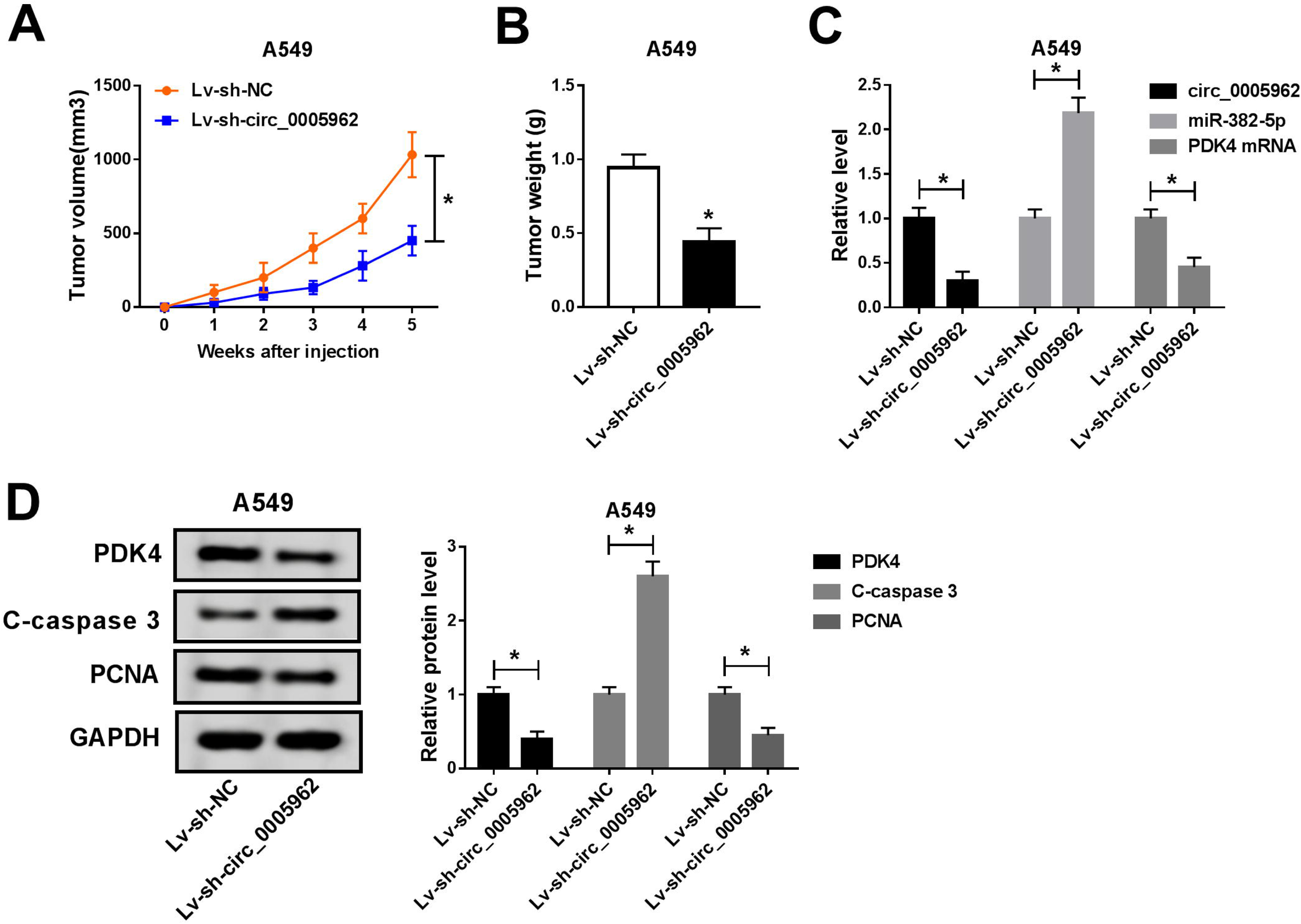
Circ_0005962 knockdown depleted tumor growth *in vivo*. (A) The tumor volume was recorded once a week, lasting 5 weeks. (B) The tumor weight was measured after 5 weeks. (C) The expression of circ_0005962, miR-382-5p and PDK was examined in excised tumor tissues by qRT-PCR. (D) The protein levels of PDK4, C-caspase 3 and PCNA were quantified by western blot. \**P* < 0.05.

## Discussion

NSCLC is a severe burden to people’s lives, and the prognosis for patients is still unsatisfactory. The responses to existing standard therapies are poor, except for the most localized cancers (Lemjabbar-Alaoui et al., 2015). Hence, more novel mechanisms of NSCLC progression need to be explored so as to develop aimed therapeutic strategies for NSCLC. Here, we investigated the role of circ_0005962 in NSCLC for the first time. Circ_0005962 was observed to be aberrantly overexpressed in NSCLC tissues and cells. The functional analysis concluded that circ_0005962 knockdown attenuated NSCLC progression *in vitro* and *in vivo*. Stepwise identification manifested that circ_0005962 could directly interact with miR-382-5p, leading to an increase of PDK4 expression, thereby contributing to the development of NSCLC. Our study illustrated the importance and carcinogenesis role of circ_0005962 in NSCLC.

CircRNAs were documented to be dysregulated in numerous human cancers and took effects on apoptosis, autophagy, chemoresistance, metastasis and glycolysis (Chi et al., 2019; Kun-Peng et al., 2018; S. Ren et al., 2019; Wei et al., 2019). Up to now, dozens of NSCLC-related circRNAs were screened and identified, such as circ_100146, CIRC-PRMT5 and circ_0026134 (Chang et al., 2019; L. Chen et al., 2019; Y. Wang, Y. Li, et al., 2019), leading to the malignant progression of NSCLC via acting as oncogenes. On the contrary, certain circRNAs, such as circPTPRA, circ_circ_00059621946 and circSMARCA5 were maintained as tumor suppressors to block NSCLC deterioration (Huang et al., 2019; Y. Wang, H. Li, et al., 2019; Wei et al., 2019). These data suggested the diverse roles of circRNAs in cancer development. Despite the fact that the role of several circRNAs was partly characterized, there were still existing circRNAs lacking functional exploration. A previous study predicted differentially expressed circRNAs in lung adenocarcinoma using Gene Expression Omnibus (GEO) dataset and found that circ_0005962 was highly expressed in lung adenocarcinoma plasma and cells (Liu et al., 2019). In view of this finding, we speculated that dysregulation of circ_0005962 might be associated with the malignant progression of NSCLC. Interestingly, we discovered that circ_0005962 knockdown inhibited cell proliferation, autophagy and glycolysis but accelerated cell apoptosis through *in vitro* analyses. Besides, circ_0005962 knockdown weakened tumor growth in nude mice *in vivo*, suggesting that circ_0005962 was an oncogene at least in NSCLC progression.

The classic action way of circRNAs is as a sponge of miRNAs. In our study, miR-382-5p was identified as a target of circ_0005962. The role of miR-382 in NSCLC gradually became clear to function as a tumor suppressor. For example, miR-382 suppressed proliferation and migration of NSCLC cells by binding to the 3’ UTR of LMO3 (D. Chen et al., 2019). Besides, miR-382 was significantly downregulated in NSCLC tissues and cells, and enrichment of miR-382 depleted cell proliferation, migration and invasion through targeting SETD8 (Chen et al., 2017). Consistent with these findings, we also detected that miR-382-5p was weakly expressed in NSCLC cells, and reintroduction of miR-382-5p inhibited proliferation, autophagy and glycolysis of NSCLC cells. Besides, miR-382-5p inhibition reversed the regulatory effects of circ_0005962 knockdown.

Considering the habitual action mode of miRNAs, we further analyzed the potential target mRNAs of miR-382-5p to establish a detailed action mechanism of circ_0005962 in NSCLC. Among the mRNAs whose levels were increased in A549 and HCC827 cells, one of the most significant was PDK4. Similarly, PDK4 has been reported to be highly expressed in cisplatin-resistant lung adenocarcinoma (Yu et al., 2018). Besides, oncogene LINC00243 contributed to proliferation and glycolysis in NSCLC by positively regulating PDK4 (Feng & Yang, 2019). Consistently, we also noticed that PDK4 was upregulated in NSCLC tissues and cell lines. Moreover, PDK4 overexpression eliminated the role of miR-382-5p reintroduction, leading to the malignant activities of NSCLC cells. These data indicated that PDK4 played a carcinogenic role in NSCLC.

Taken together, the expression of circ_0005962 was increased in NSCLC tissues and cell lines. Knockdown of LINC00243 blocked cell proliferation, autophagy and glycolysis but accelerated cell apoptosis *in vitro* and weakened tumor growth *in vivo*. Besides, circ_0005962 was a sponge of miR-382-5p, and PDK4 was a target of miR-382-5p. Circ_0005962 functioned in NSCLC progression by inducing PDK4 through sponging miR-382-5p. Our results not only corroborate the role of circ_0005962 in NSCLC *in vitro* and *in vivo* but also provide the circ_0005962/miR-382-5p/PDK4 regulatory axis, which may be promising to develop novel therapeutic approaches for NSCLC.

## Acknowledgement

None

## Author contributions

Conceptualization: Z.-H.Z.; Methodology: Z.-X.S., R.-B.C., X.-R.P.,; Software: X.-R.P., B.-X., L.-X.; Validation: Z.-H.Z., Z.-X.S.; Formal analysis: G.-F.Z.; Investigation: Z.-X.S., R.-B.C., X.-R.P.; Resources: B.-X., L.-X.; Writing - original draft: Z.-H.Z.

